# AARS Online: a collaborative database on the structure, function, and evolution of the aminoacyl-tRNA synthetases

**DOI:** 10.1101/2024.05.15.594223

**Authors:** Jordan Douglas, Haissi Cui, John J. Perona, Oscar Vargas-Rodriguez, Henna Tyynismaa, Claudia Alvarez Carreño, Jiqiang Ling, Lluís Ribas-de-Pouplana, Xiang-Lei Yang, Michael Ibba, Hubert Becker, Frédéric Fischer, Marie Sissler, Charles W. Carter, Peter R. Wills

## Abstract

The aminoacyl-tRNA synthetases (aaRS) are a large group of enzymes that implement the genetic code in all known biological systems. They attach amino acids to their cognate tRNAs, moonlight in various non-translational activities, and are linked to many genetic disorders. The aaRS have a subtle ontology characterized by structural and functional idiosyncrasies that vary from organism to organism, and protein to protein. Across the tree of life, the twenty-two coded amino acids are handled by sixteen evolutionary Families of Class I aaRS and twenty-one Families of Class II aaRS. We introduce AARS Online, an interactive Wikipedia-like tool curated by an international consortium of field experts. This platform systematizes existing knowledge about the aaRS by showcasing a taxonomically diverse selection of aaRS sequences and structures. Through its graphical user interface, AARS Online facilitates a seamless exploration between protein sequence and structure, providing a friendly introduction to the material for non-experts and a useful resource for experts. Curated multiple sequence alignments can be extracted for downstream analyses. Accessible at www.aars.online, AARS Online is a free resource to delve into the world of the aaRS.

## Introduction

The aminoacyl-tRNA synthetases (aaRS) are a diverse group of enzymes that attach amino acids to their cognate tRNAs [1]. These enzymes implement the genetic code in all known organisms, in all domains of life – the bacteria, archaea, and eukarya – as well as mitochondria and chloroplasts [2], and are also expressed by certain viruses [3]. Due to their fundamental roles in building and managing living cells, aaRS mutants are linked to many diseases in higher organisms [4], where they carry out a wide range of additional functions that are not directly related to protein synthesis [5].

Aminoacylation is a two-step reaction powered by adenosine triphosphate (ATP) [6]. Adenosine monophosphate (AMP) and pyrophosphate (PP) are released as byproducts of the reaction:

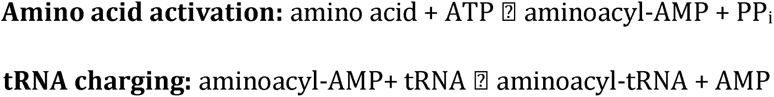

These two reactions are carried out by the catalytic domain, which exists in two distinct evolutionary forms: Class I and Class II. The Class I catalytic domain is a Rossmann fold, with four parallel -strands, and Class II an anti-parallel -sheet with six strands. Nine of the 22 coded amino acids are supplied to the ribosomal machinery from tRNAs charged exclusively by Class I enzymes, eleven by Class II, and the remaining two amino acids, lysine [7] and cysteine [8], can be rendered variously from the products of either Class I or II analogs. The second reaction, tRNA charging, is facilitated through recognition of the cognate tRNA by numerous structural elements of the catalytic domain [9], and one or more additional domains that bind to the anticodon stem [10], the D-loop [11], the variable arm [12], or other parts of the tRNA molecule. Mis-activated and mis-charged amino acids can be expelled from the reaction pathway through editing activity, which operates at the pre-transfer and post-transfer level, respectively [1]. Editing is necessary when varying amino acid types cannot be accurately distinguished by the active sites of the enzymes [6]. Some aaRS possess an additional domain that catalyzes post-transfer editing [13], [14].

In most cases, each aaRS supplies just a single type of amino acid onto the growing peptide, and that amino acid is specified in the naming of the enzyme, for example alanyl-tRNA synthetase (AlaRS) attaches alanine to tRNA. However, certain aaRS can supply an additional amino acid or provide a building block, which is then elaborated into the amino acid corresponding to the tRNA. First, the metazoan glutamyl-prolyl-tRNA synthetase (EPRS) is a fusion of the catalytic domains from GluRS and ProRS connected by a linker and hence supplies both amino acids to their respective tRNA. Second, the original amino acid can be changed after its attachment to tRNA before reaching the ribosome. This is the case for the non-discriminating aspartyl- and glutamyl-tRNA synthetases (AsxRS and GlxRS), which attach Asp to tRNA and Glu to tRNA, respectively [15], [16]. The rare GluGlnRS attaches Glu to tRNA in a discriminating manner, representing an ancestral midpoint between the discriminating and non-discriminating modes [17], [18]. Likewise, O-phosphoseryl-tRNA synthetase (SepRS) supplies cysteine for organisms that lack CysRS [8], and SerRS charges serine to both tRNA^Ser^ and tRNA^Sec^ [19].

All human aaRS are encoded by nuclear genes, and all of them have been linked to genetic diseases [4], [20]–[24]. 19 aaRS enzymes remain in the cytosol (EPRS doubles up) and 19 are relocated to mitochondria (no mitochondrial GlnRS) [20]. The cytosolic and mitochondrial aaRS are encoded by separate genes, with the exception of GlyRS and LysRS, which are shared by both compartments through dual targeting signals [25]. Each protein is encoded by a single gene with the exception of the cytosolic PheRS protein, whose and subunits are expressed by two genes. Therefore humans have a total of 37 aaRS genes and 36 proteins (splice variants excluded). Genes that produce cytosolic and mitochondrial aaRS are conventionally named XARS1 and XARS2 respectively, e.g., CARS1 and CARS2 encode cytosolic and mitochondrial cysteinyl-tRNA synthetase, and the latter often resemble bacterial aaRS, due to the endosymbiotic origin of mitochondria [26]. The dominant cytosolic aaRS mutations typically affect the peripheral nervous system (e.g., Charcot-Marie-Tooth disease [27]) while the recessive mutations in cytosolic aaRS tend to affect a large range of organs and are often accompanied with developmental delays (e.g., microcephaly [28]) [29]. Mitochondrial aaRS mutations typically affect organs of high metabolic demand, such as the brain and heart, as well as other tissues (e.g., leukodystrophy [30]) [22], [31], [32]. In some cases, mutations in the same aaRS result in strikingly different disease phenotypes, but in most cases the causal mutations of a disorder can occur at many positions in the respective aaRS, even across different domains [21].

Protein domains in general display a tendency to rearrange, duplicate, and exchange as functional modules on evolutionary timescales [33]–[35], and the aaRS are no exception to this phenomenon. Several aaRS have distinct isoforms that arose from gene fusion or fission events, such as the metazoan EPRS [36] and the parasitic TyrRS [37], which each have two catalytic domains, and the bacterial GlyRS, which usually consists of and subunits _2 2_, but occurs as an _2_ fusion in some bacteria and chloroplasts [38], [39]. Fused and fissed isoforms also exist for AlaRS [40], PheRS [41], and PylRS [42]. These relatively recent gene rearrangement events shed light on deeper phylogenetic incongruences across the respective Class. Namely, LysRS-I and GluRS have paralogous anticodon binding domains, but their catalytic domains are more divergent (**Fig. 1**). And the anticodon binding domain of HisRS is related to ProRS and ThrRS, while its catalytic domain is more similar to PheRS (**Fig. 2**). These “domain hopping” events have led to conflicting phylogenetic interpretations of LysRS-I and HisRS in the past [9], [43]–[46]. Similarly, AlaX proteins remove mischarged amino acids from tRNA, and also appear as a C-terminal domain of AlaRS and an N-terminal domain of ThrRS [47]–[49], while various eukaryotic extensions, such as the GST-like and WHEP domains, also occur elsewhere in the proteome [50].

**Fig 1.**
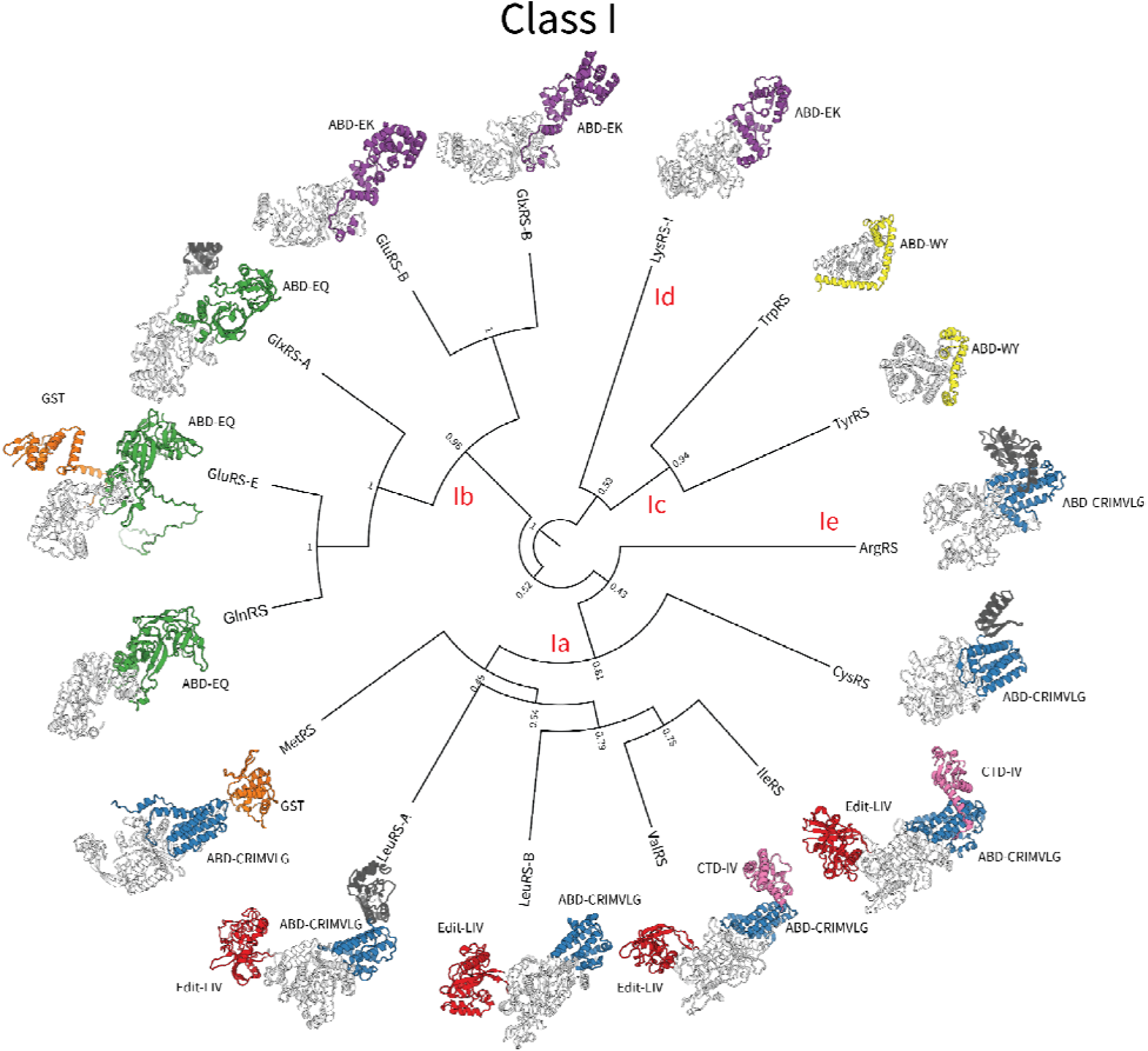
An inferred phylo geny of the Class I catalytic domain [9]. Protein structures are colored by domain Superfamily - catalytic domain: light gray, ABD-EK: purple, ABD-EQ: green, ABD-CRIMVLG: blue, ABD-WY: yellow, Edit-LIV: red, GST: orange, all other domains: dark gray. Branch lengths are proportional to amino acid substitutions per site, and internal nodes are labeled by clade posterior support. Please refer to Table 2 for domain Superfamilies.

**Fig 2.**
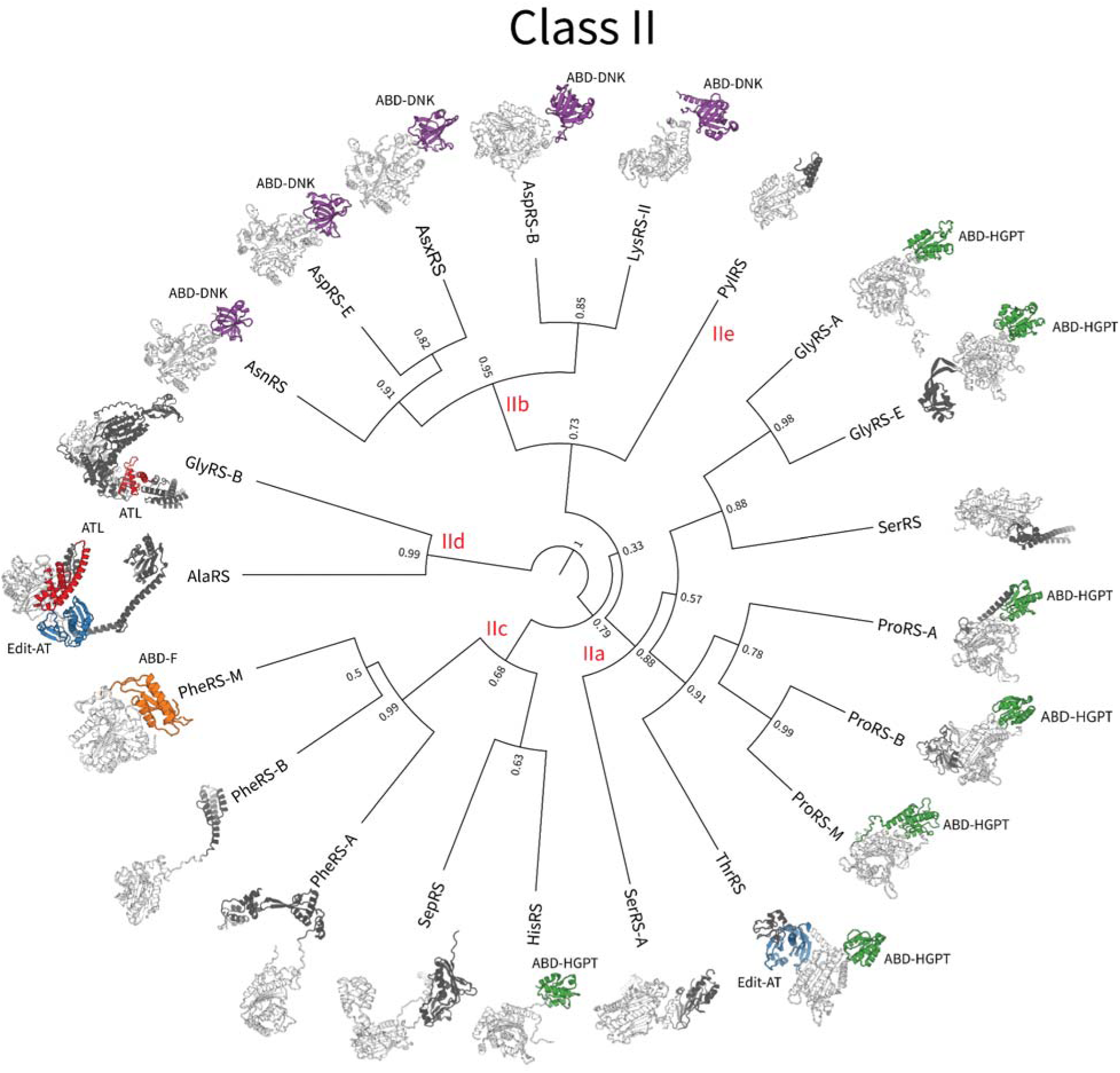
An inferred phylogeny of the Class II catalytic domain [9]. Protein structures are colored by domain Superfamily - catalytic domain: light gray, ABD-DNK: purple, ABD-HGPT: green, ABD-F: orange, Edit-AT: blue, ATL: red, all other domains: dark gray. Branch lengths are proportional to amino acid substitutions per site, and internal nodes are labeled by clade posterior support. ABD-DNK also occurs in some of the Class I aaRS in the form of EMAP. Note that the chains of PheRS were omitted from this phylogeny, as their catalytic domain paralog has diverged from the rest of the Superfamily, and the chains of GluRS-B were omitted as the y do not possess a paralog of the catalytic domain at all (the GlyRS-B structure displayed is a fusion of the two chains). Please refer to Table 2 for domain Superfamilies.

Accordingly, each aaRS is herein classified at three taxonomic levels: Class, Subclass, and Family, based on its catalytic domain (**Table 1**), which is the only domain common to the aaRS [51]. First, there are two catalytic domain *Classes*: Class I and Class II. All members of a Class share common ancestry and are therefore protein Superfamilies under the CATH and SCOP frameworks [52], [53]. Second, aaRS are further divided into *Subclasses*, based on their sequence and structural similarities. However, there is no general consensus on Subclass assignment [1], [9], [43]–[46], [54]. These discrepancies arise from a combination of faded phylogenetic signals (these proteins are over three billion years old), the use of varying phylogenetic methodologies and datasets, and also depend on whether the classification was based on the full-length protein or the catalytic domain only. In any case, members of a Subclass tend to recognize amino acids that have similar properties, for instance the branched chain amino acids (Leu, Ile, and Val) are all supplied by the enzymes of Subclass *Ia*, and large aromatic side chains (Trp and Tyr) by Subclass *Ic*. Third, following the nomenclature by Douglas et al. 2024 [9], a *Family* is a set of catalytic domains that have the same aminoacylation activity, are monophyletic (or are monophyletic with a second Family contained within them), and cannot be further distinguished by an insertion or deletion of at least 50 amino acids. In the case of multiple Families sharing the same function, they are named after their predominant taxonomic group: ‘A’ for archaeal-like, ‘B’ for bacterial-like, ‘E’ for eukaryote-like, and ‘M’ for mitochondrial-like. As the aaRS are often dual targeted to mitochondria and chloroplasts [2], with information on chloroplastic aaRS being scarce, the ‘M’ moniker does not discriminate between either compartment. As a tiebreaker, the ‘A’ label takes precedence over ‘E’, as eukaryotes can be regarded as specialized forms of archaea [55]. As shown in **Table 1**, there are many instances of common aminoacylation functions that fall under multiple Families, such as glycyl-tRNA synthetase, which exists as bacterial-like, archaeal-like, and eukaryote-like forms (GlyRS-B, GlyRS-A, and GlyRS-E), and the two forms of leucyl-tRNA synthetase (LeuRS-A and LeuRS-B) whose editing domains are nested in distinct regions of the catalytic domain [56].

**Table 1.**
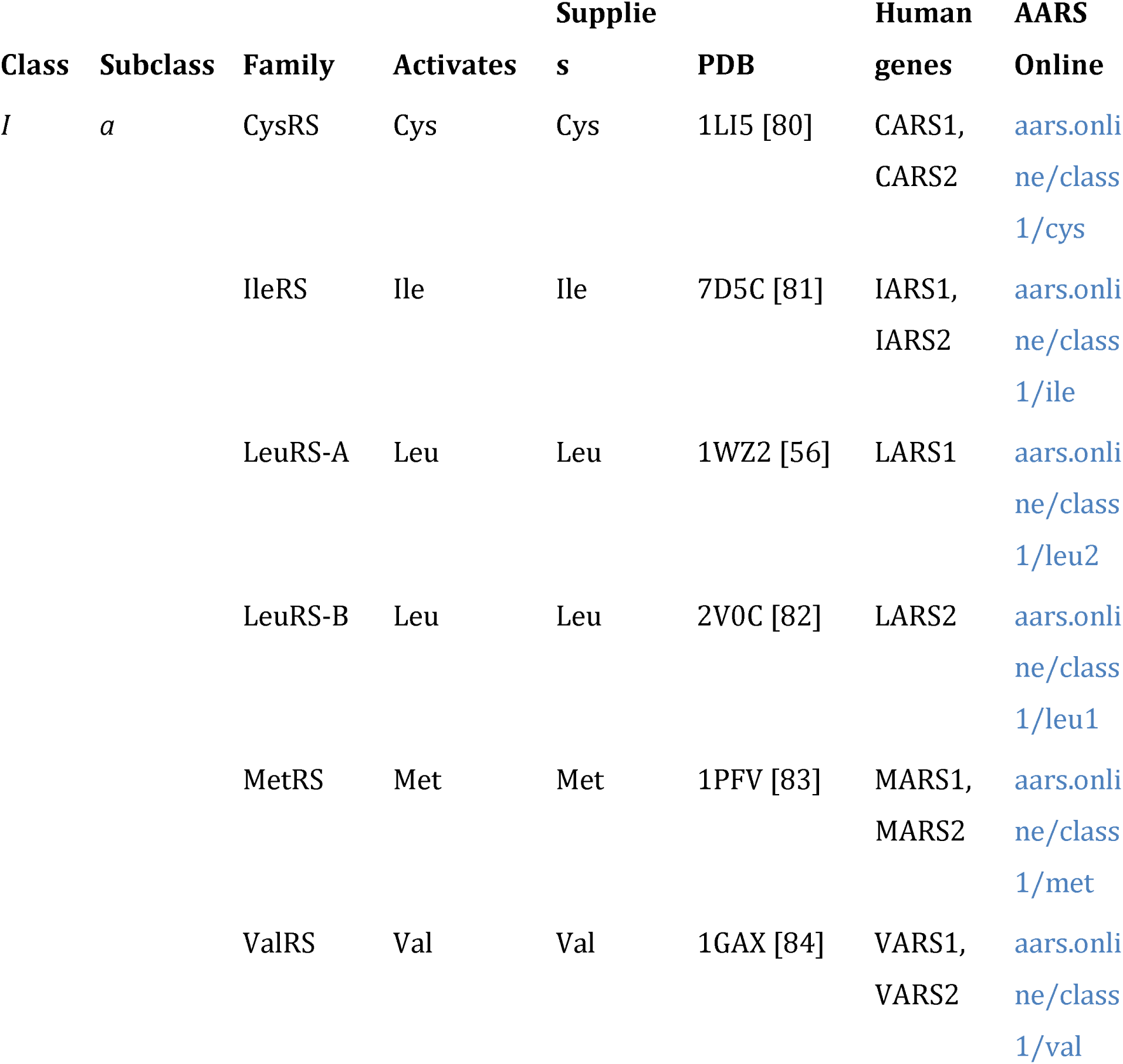
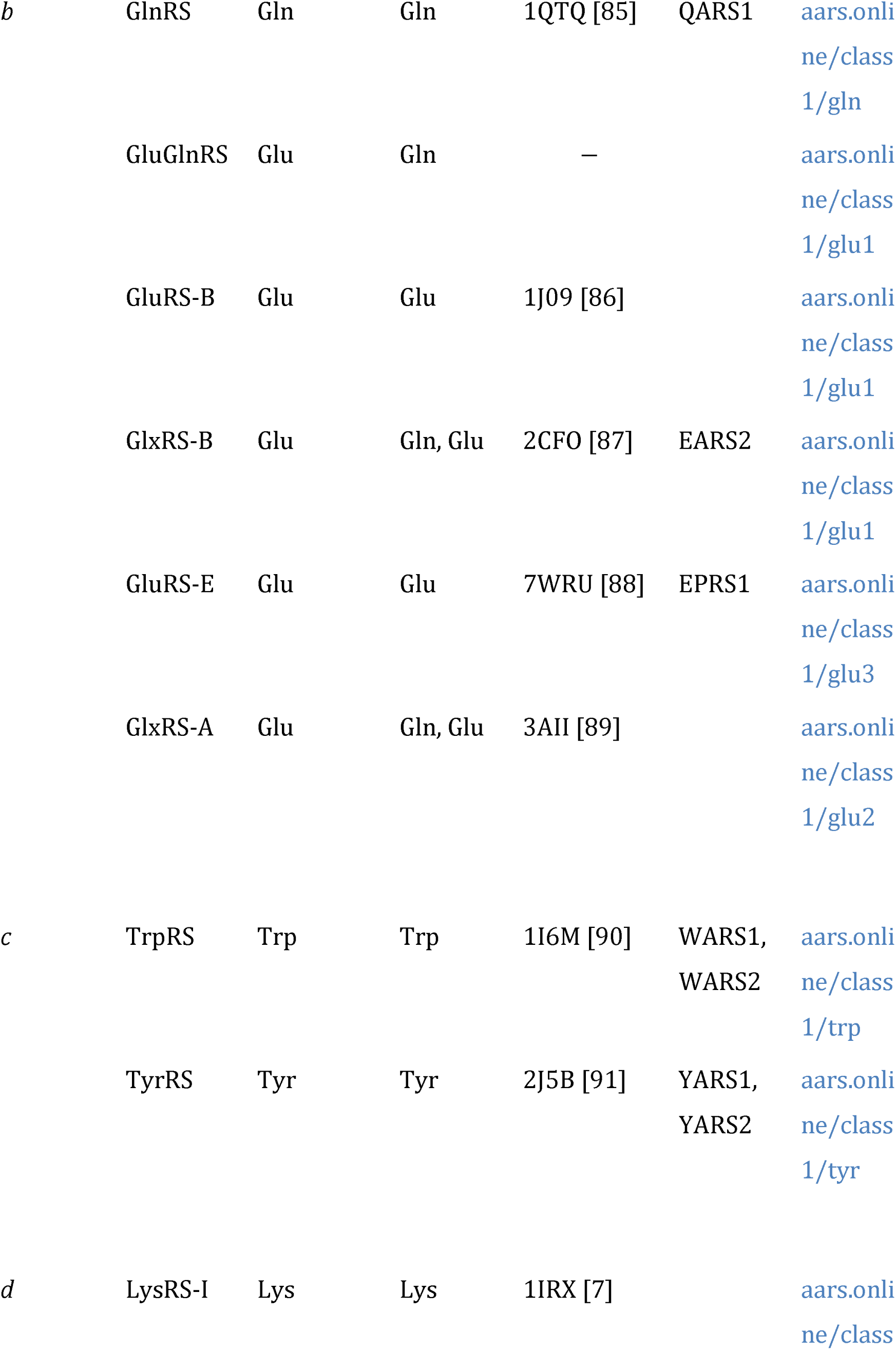
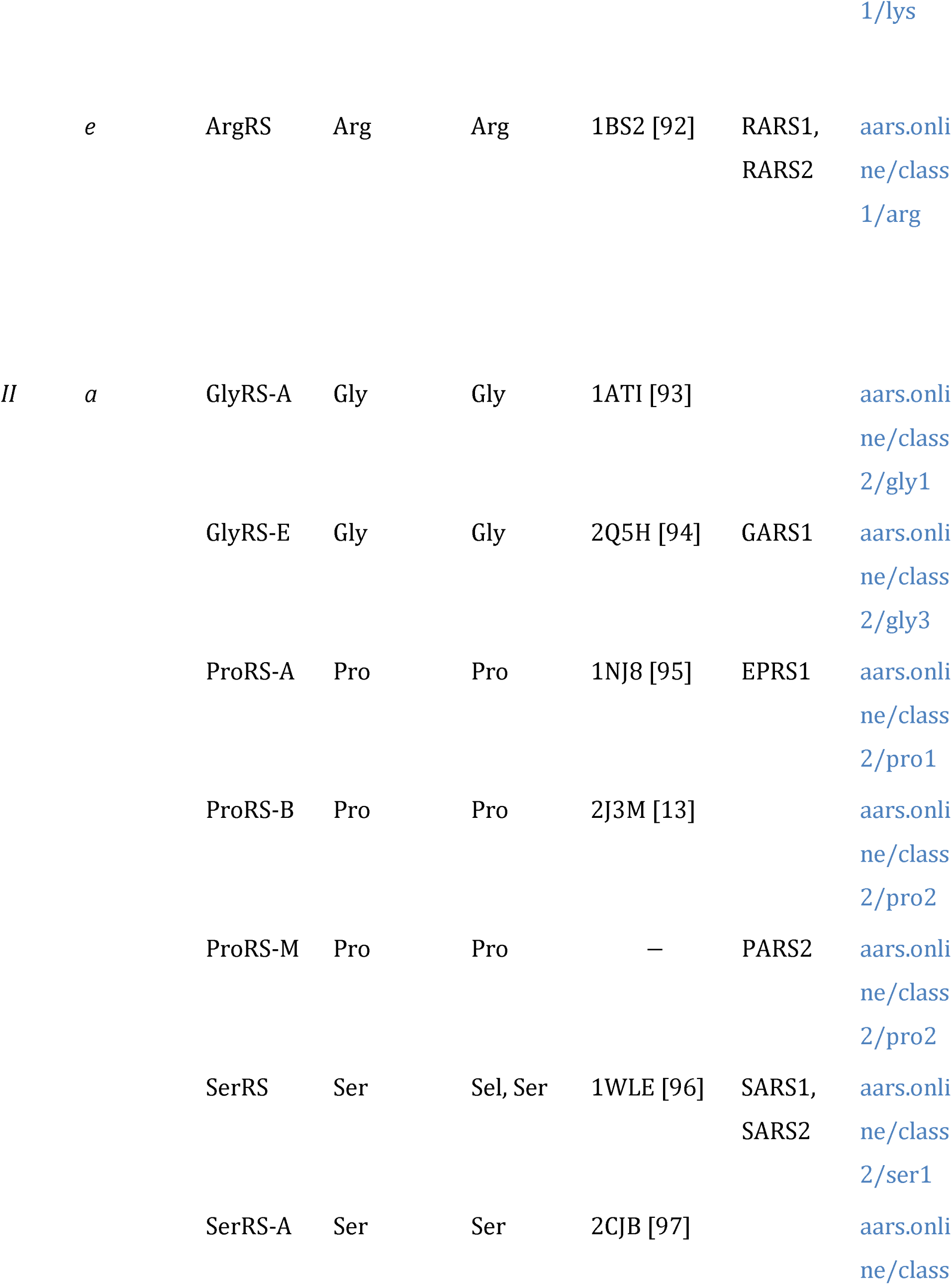
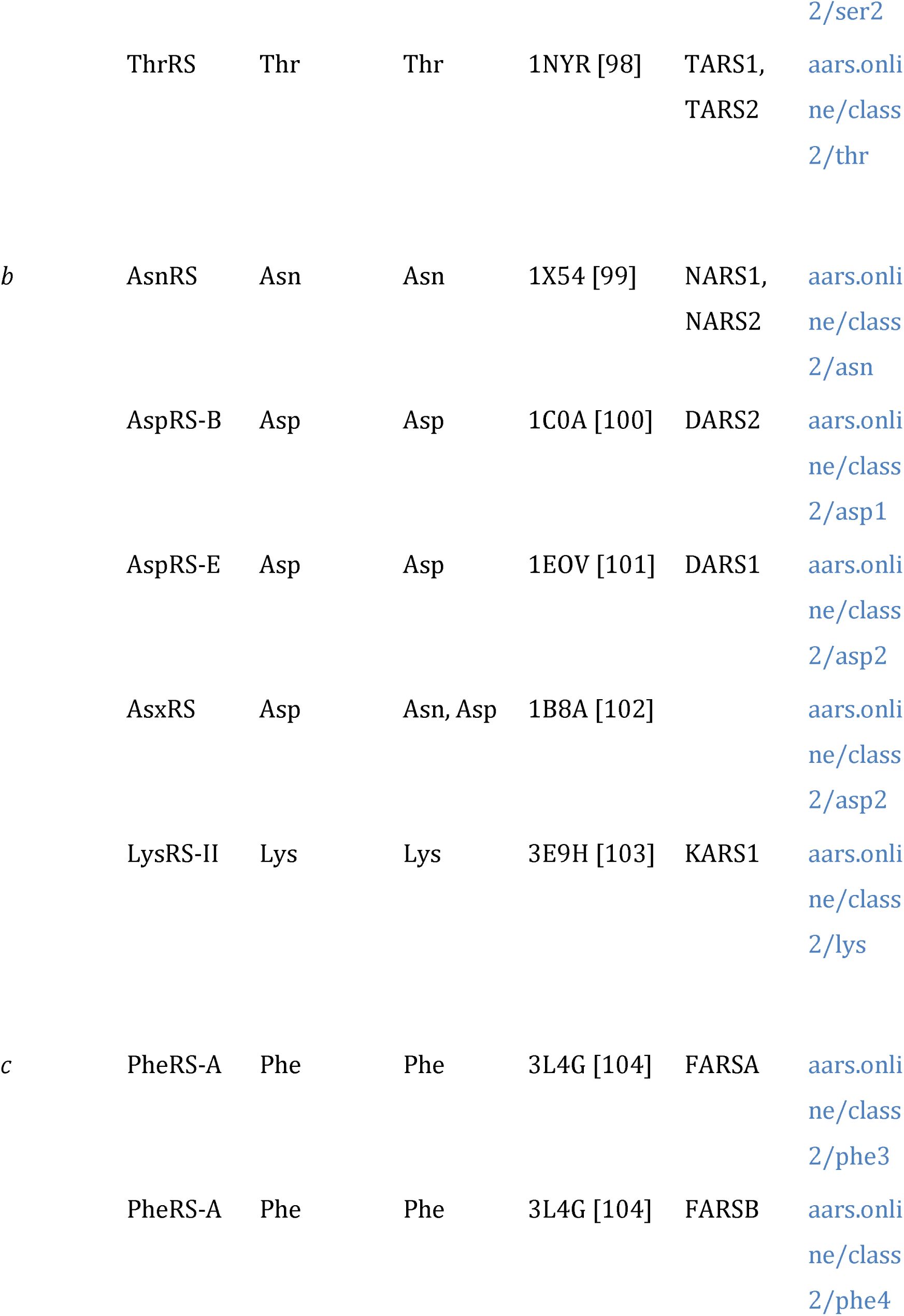
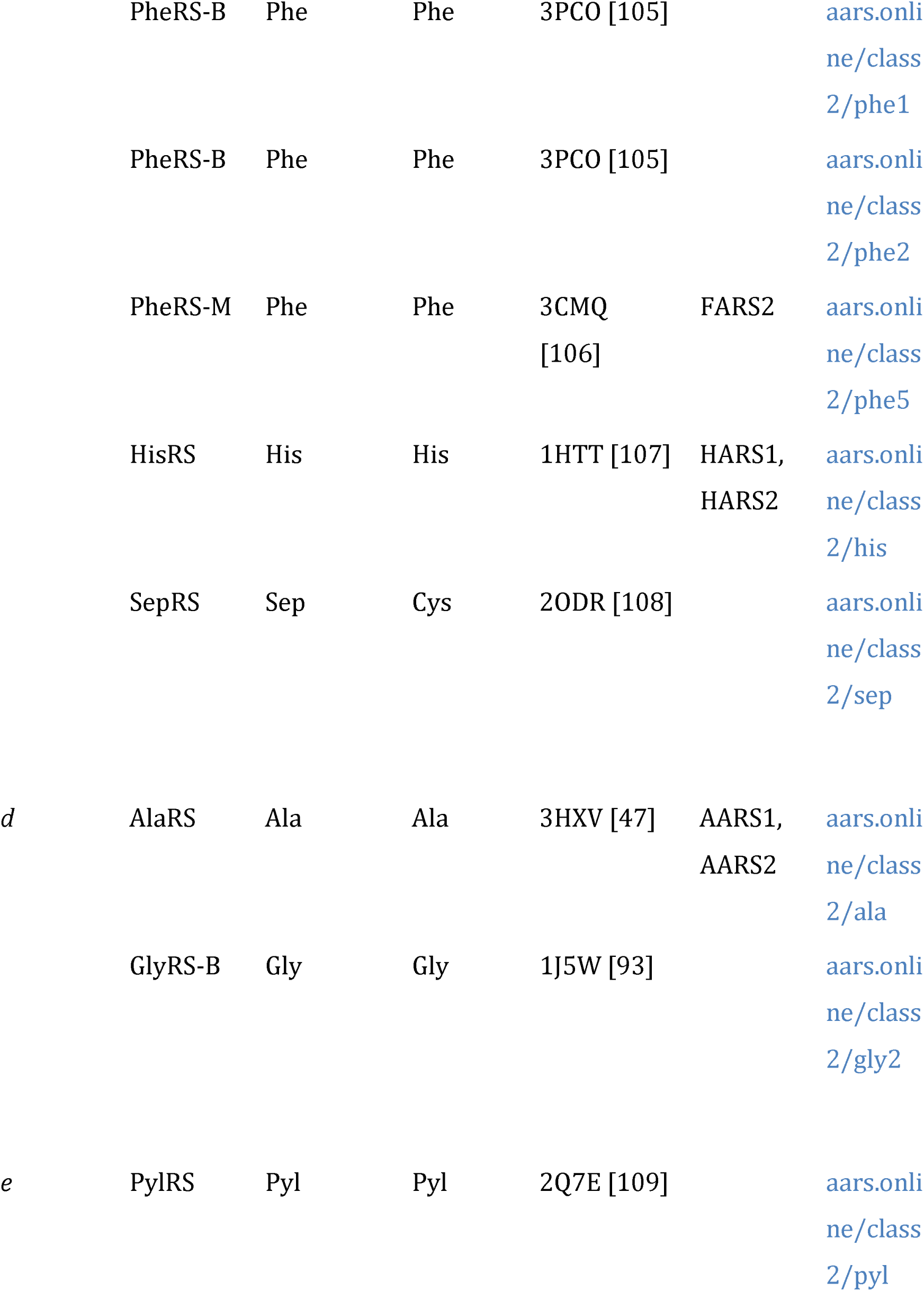
Summary of aaRS Families. Where available, an example of a solved structure is specified by its Protein Data Bank (PDB) code. The tetrameric PheRS contains paralogs of the Class II catalytic domain in the and chains, however catalysis is confined to the subunit [41]. In contrast, the subunit of GlyRS-B does not have a paralog of the catalytic domain, and hence does not have a web page.

Remarkably, the evolutionary history of the aaRS traces back to primordial life before the last universal common ancestor [57]. Under the nucleopeptide world hypothesis (see reviews: [58], [59]), the genetic code originated in an environment governed by the RNA catalysis of peptide synthesis, and the peptide catalysis of RNA synthesis. The seemingly-unrelated Class I and II aaRS plausibly originated as proteins that co-docked tRNA molecules from opposing sides [43], and encoded by opposing strands of a bidirectional gene [60], [61], which would have served as a central integrating role between the RNA and peptide populations [61], [62]. These small, primitive aaRS, known as “urzymes” [63]–[67], would have been promiscuous enzymes that may have recognized small, primitive forms of tRNA, known as “minihelices” [68]–[71]. Nucleopeptide world is a direct challenge to the classical RNA world hypothesis (see reviews: [59], [72]), which postulates that genetic coding originated in a world governed by self-replicating RNA catalysts, including ribozymal aminoacyl-tRNA synthetases [73] that were later supplanted by the proteinaceous forms we know today. Many laboratories have successfully engineered ribozymes that carry out one of the two aaRS reactions, but not both [74]–[76].

There are representative solved structures for the majority of known aaRS Families (**Table 1**). However, given the extensive resources needed to produce just one structure, those available are often sourced from model organisms, or are chosen for their medical/economic relevance or their ability to crystallize, and are thus far from a representative sample of the biosphere’s full diversity. For instance, the three most common source organisms on the entire Protein Data Bank are all eukaryotic, all multicellular, and all vertebrates – humans, cattle, and chickens (May 2024). Likewise, a disproportionately large number of solved aaRS structures were sourced from *Escherichia coli*, *Saccharomyces cerevisiae*, and *Thermus thermophilus*. The advent of AlphaFold [77] means that protein structures can be accurately predicted, moreso when the protein has solved homologs, and therefore these taxonomic sampling biases can be addressed. While AlphaFold models are still less reliable than experimentally determined structures [78], [79], they can be useful for generating experimentally testable hypotheses when considered in conjunction with other lines of evidence.

Existing knowledge about the aaRS is vast, and has been assembled from a range of academic perspectives, including enzymology, structural biology, cell biology, biomedicine, and phylogenetics. Navigating the plethora of aaRS Families across these diverse academic fields can be challenging. Here we introduce AARS Online – a platform that systematizes aaRS knowledge across the tree of life, built by, and for the use of, the aaRS community. At its core, AARS Online is a user-friendly Wikipedia-like tool for interactively displaying aaRS structures, sequences, and evolutionary information side-by-side. The platform is open-source and hosted on GitHub (https://github.com/aarsonline/aarsonline.github.io), promoting users to edit and contribute content, ensuring that material remains up-to-date. This resource can be accessed at https://www.aars.online.

## Materials and Methods

### Sequence and structure alignment

We selected a digestible number of sequences and structures for each aaRS Family - between five and twenty five. These samples consist of solved and predicted protein structures, sourced from both model organisms and the rest of the biosphere. We used the following inclusion criteria for selecting sequences/structures. First, we included experimentally solved aaRS structures from the Protein Data Bank. Second, we extracted additional annotated aaRS sequences from GenBank and predicted their monomeric structures using AlphaFold v2.3.0 [77]. Most of these AlphaFold models were generated by the New Zealand eScience Institute cluster, and others by ColabFold [110]. We ensured that the aaRS from the following model organisms were included (as either solved or AlphaFold structures): *Escherichia coli*, *Homo sapiens* (cytosolic and mitochondrial), and *Saccharomyces cerevisiae* (cytosolic only). We randomly sampled other sequences/structures for inclusion in a taxonomically-representative fashion, such that each aaRS Family had a minimum of four samples from four phyla, and where possible, up to eight bacterial phyla, four archaeal phyla, four eukaryotic phyla, and one viral phylum, plus two organellar (mitochondrial or chloroplast) samples from two distinct eukaryotic phyla.

Protein three dimensional structures were displayed using PV^1^. Secondary structures were defined using DSSP v3.0.0 [111], [112]. Pairwise structural alignments were generated by DeepAlign [113]. Per-Family multiple sequence alignments were generated by first aligning the structures with 3DCOMB [114], followed by a refinement algorithm that realigned contiguous regions of at least three sites lacking secondary structure, using ClustalW [115] based on primary structure [9]. As existing structural alignment tools were not always reliable at delineating homologous insertions, alignments were then adjusted manually.

Realignment and adjustment was especially important for the Class I KMSKS motif, whose structure is highly flexible and yet its sequence is strongly conserved.

Most of the protein structures displayed in **Fig. 1** and **Fig. 2** are experimental structures, whose PDB codes are specified in **Table 1**. However, in order to capture additional domains that were missing from experimental structures, we used AlphaFold structures for the following Families: CysRS (species: *Aciduliprofundum boonei*; GenBank: 8827372), MetRS (*Scheffersomyces stipitis*; 4836672), GlyRS-B (*Chlamydia pneumoniae*; 45051002), GluRS-E (*Cyathus striatus*; KAF9013924), and GlxRS-A (*Sulfolobus islandicus*; 15298417). Cognate tRNA secondary structures were extracted from tRNAViz [116].

### Phylogenetics

The catalytic domain phylogenies in **Fig. 1** and **2** are from Douglas et al. 2024 [9] and were displayed using FigTree^2^. The remaining phylogenetic analyses presented on AARS Online were performed using BEAST v2.7.6 [117]. All phylogenies presented on AARS Online were inferred under the optimized relaxed clock (ORC v1.2.0) [118], the OBAMA substitution model (OBAMA v1.1.1) [119], and the Yule Skyline tree prior (BICEPS v1.1.1) [120]. Trees were summarized using the CCD0 tree [121]. Markov chain Monte Carlo was run until the effective sample sizes of all parameters exceeded 200, diagnosed using Tracer v1.6 [122]. The cross-Family catalytic domain phylogenies were constructed using the common elements of the catalytic domain of either Class. The OBAMA method selected the WAG substitution model [123] for Class I (with 100% posterior support), and the LG model [124] for Class II (also with 100% support).

### Web content

AARS Online is hosted on GitHub and was coded entirely in JavaScript. Curators are able to edit the web page introductions by editing markdown files on GitHub, which automatically deploys any changes made. Users are able to submit pull requests to make changes, which will be deployed pending acceptance by a curator. All computation and database indexing is performed client-side, which avoids congestion at a back-end server.

## Results and Discussion

### Sequence and structural data

In total, AARS Online includes 545 aaRS structures: 68 were extracted from the Protein Data Bank and 477 were generated using AlphaFold. This taxonomically diverse sample was sourced from 49 phyla of the tree of life: 228 structures are bacterial (from 17 bacterial phyla), 113 archaeal (8 phyla), 137 from the eukaryotic cytosol (17 phyla), 60 from the eukaryotic organelles (14 phyla), and 7 viral (single phylum). This viral phylum corresponds to the giant viruses, which have collected several aaRS in their abnormally large genome [3]. Although taxonomy is dynamic, especially in the case of archaea, and many life forms remain undiscovered or unsequenced, much of the aaRS diversity across the tree of life has likely been captured by this sample.

The reliability of the AlphaFold models can be evaluated on AARS Online by displaying the per-site pLDDT scores on the 3D protein structures. The pLDDT scores were generally quite large, providing some level of confidence in the structural models. All aaRS Families had a median score of at least 94% for the catalytic domain and at least 91% for the full-length protein. The chain of PheRS-A was the lowest scoring aaRS Family, whose catalytic domains had a median score of 95%, but the full-length proteins were slightly lower at 91% due to the flexible linker between the two domains. The generally high scores reflect the abundance of experimentally solved aaRS structures, which were used to train AlphaFold. Low scoring regions of the aaRS generally correspond to domain linkers, and flexible or disordered regions [9], such as the KMSKS motif of Class I [125] and the flexible loop found on the surface of CysRS [80].

### Website layout

In AARS Online, there is one web page per Family (see **Table 1**), and one per aaRS domain Superfamily that is represented by more than one Family (see **Table 2**). The pairwise alignment web pages provide comparisons between Families. These web pages contain a pairwise multiple sequence/structure alignment between every pair of catalytic domain Families from the same Class. Unlike the Family and domain pages, the content on the pairwise pages was automatically generated, and they were not treated to any level of manual curation or documentation.

**Table 2.**
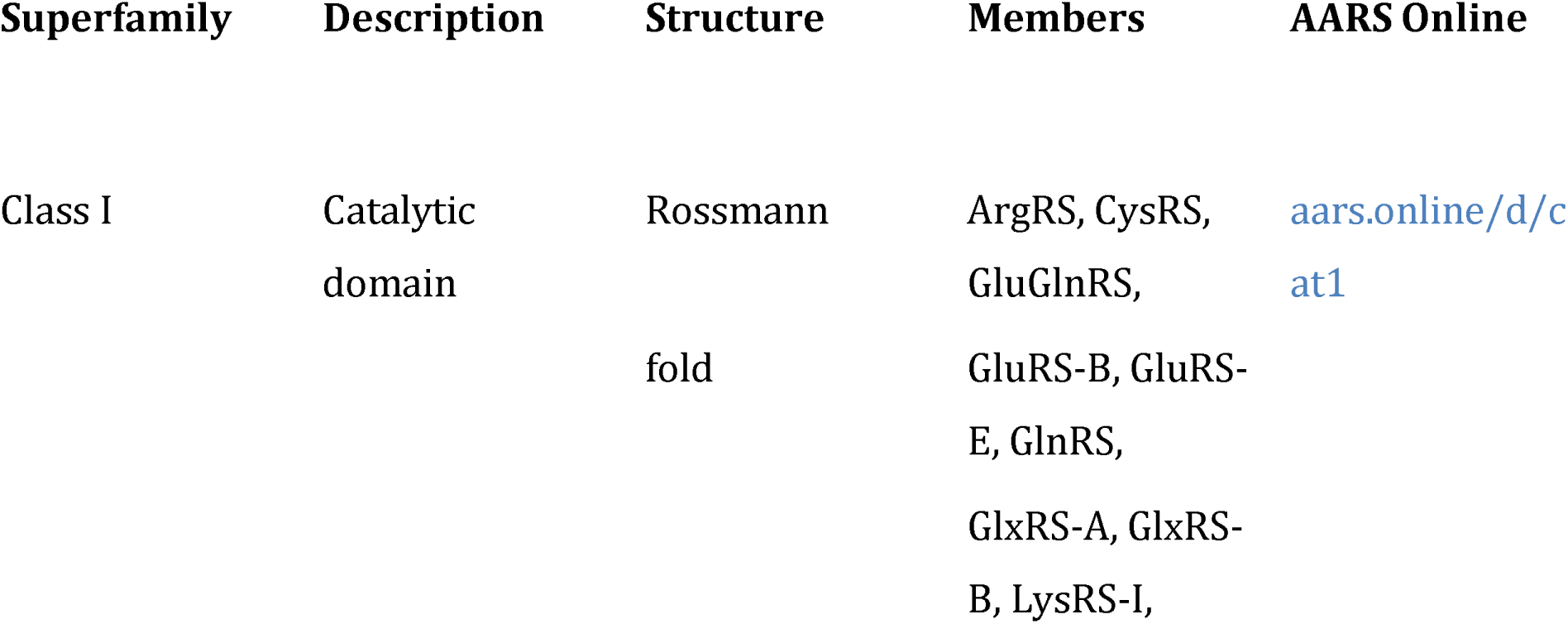
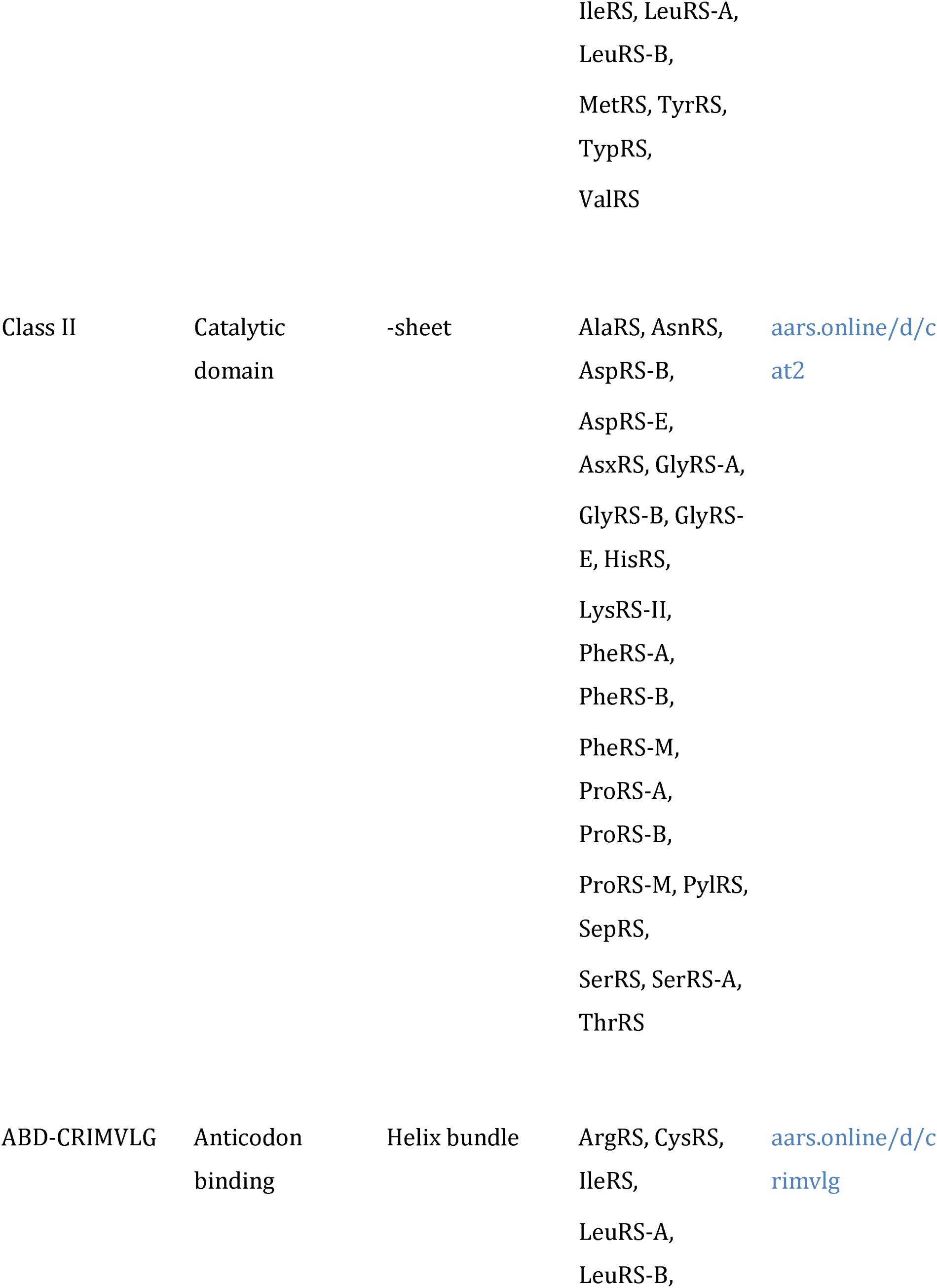
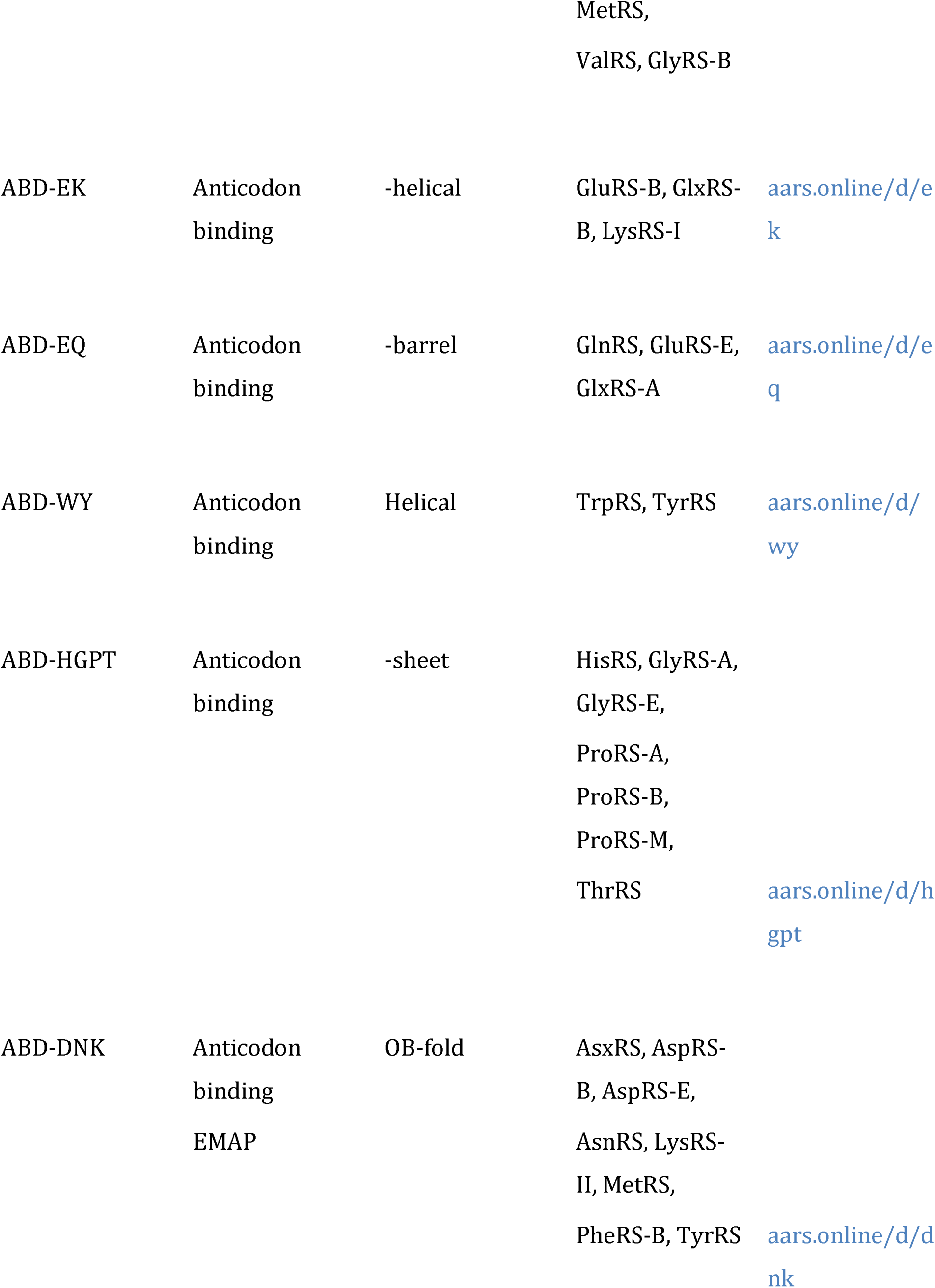
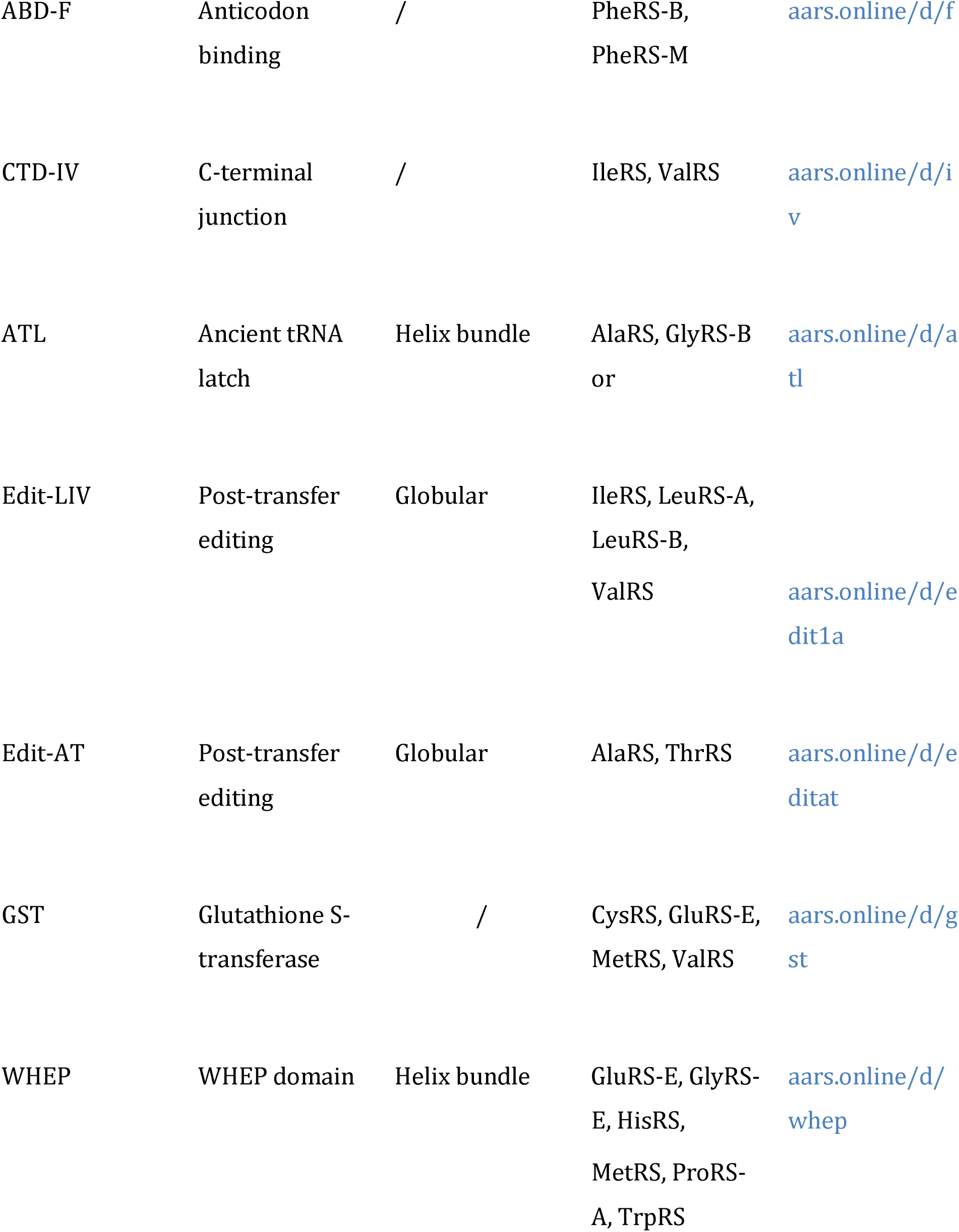
aaRS domain Superfamilies. Shown are domains that occur in at least two of the Families presented in Table 1. Each row in this table is accompanied by a web page in AARS Online. The ATL domain occurs in the chain of most _2 2_ GlyRS-B proteins, and in the fused _2_ protein in some bacteria and chloroplasts [133].

Each Family and domain web page begins with an introduction that provides a fully referenced mini-review of the system (**Fig. 3**). These introductions cover the following topics where applicable: structure and function, editing, non-translational functions, clinical significance, and tRNA. The clinical significance sections discuss, where appropriate, the role that the cytosolic and mitochondrial aaRS genes play in human disease, and their role in antibiotic resistance. As AARS Online is hosted on GitHub, changes to the database and documentation are automatically tracked. To promote user engagement, each web page features a hyperlink to GitHub, where users are encouraged to initiate discussion or provide corrections and updates by submitting pull requests.

**Fig 3.**
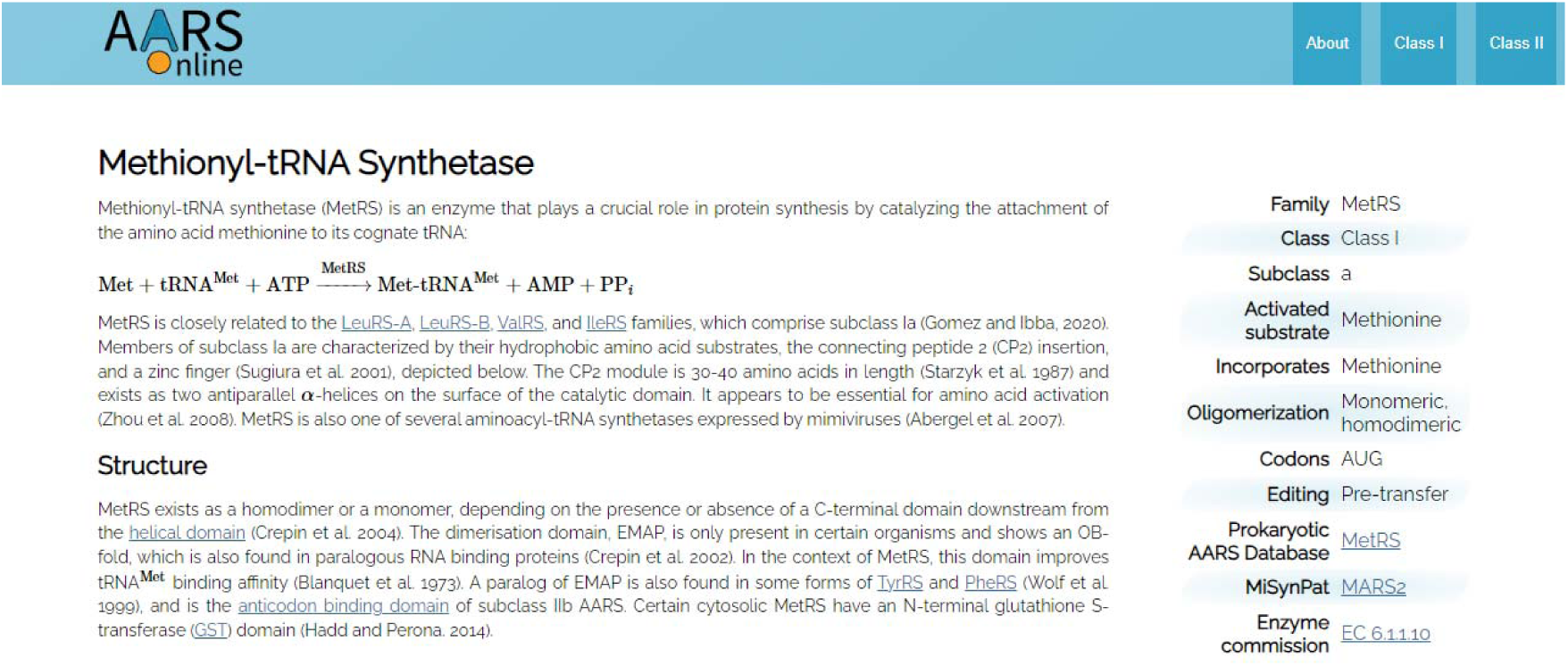
. Screenshot of the introduction on the MetRS web page.

Further down the page are annotated multiple sequence and structure alignments, a phylogenetic tree describing the relationships between proteins, and cognate tRNA secondary structures (with identity elements indicated, based on Giegé and Eriani 2023 [126]). The aaRS structures displayed are a combination of AlphaFold structures, subject to representative sampling from all domains of life (including organelles and viruses), and experimentally solved structures, subject to their availability. Sites in the multiple sequence alignment can be selected, and the corresponding areas on the protein structure(s) are also highlighted, allowing for a seamless exploration between sequence and structure (**Fig. 4**). Primary and secondary structure alignments are displayed - the former is represented by the 20 canonical amino acid characters (plus gaps), and the latter by the eight DSSP secondary structure characters (plus gaps). These eight characters are: E - extended -strand; H helix; G - 310 helix; I - p-helix; B - b-bridge; S - bend; T - H-bonded turn; and N - other/unclassified [111], [112]. These sequence and structural alignments can be downloaded and further interpreted with other software.

**Fig 4.**
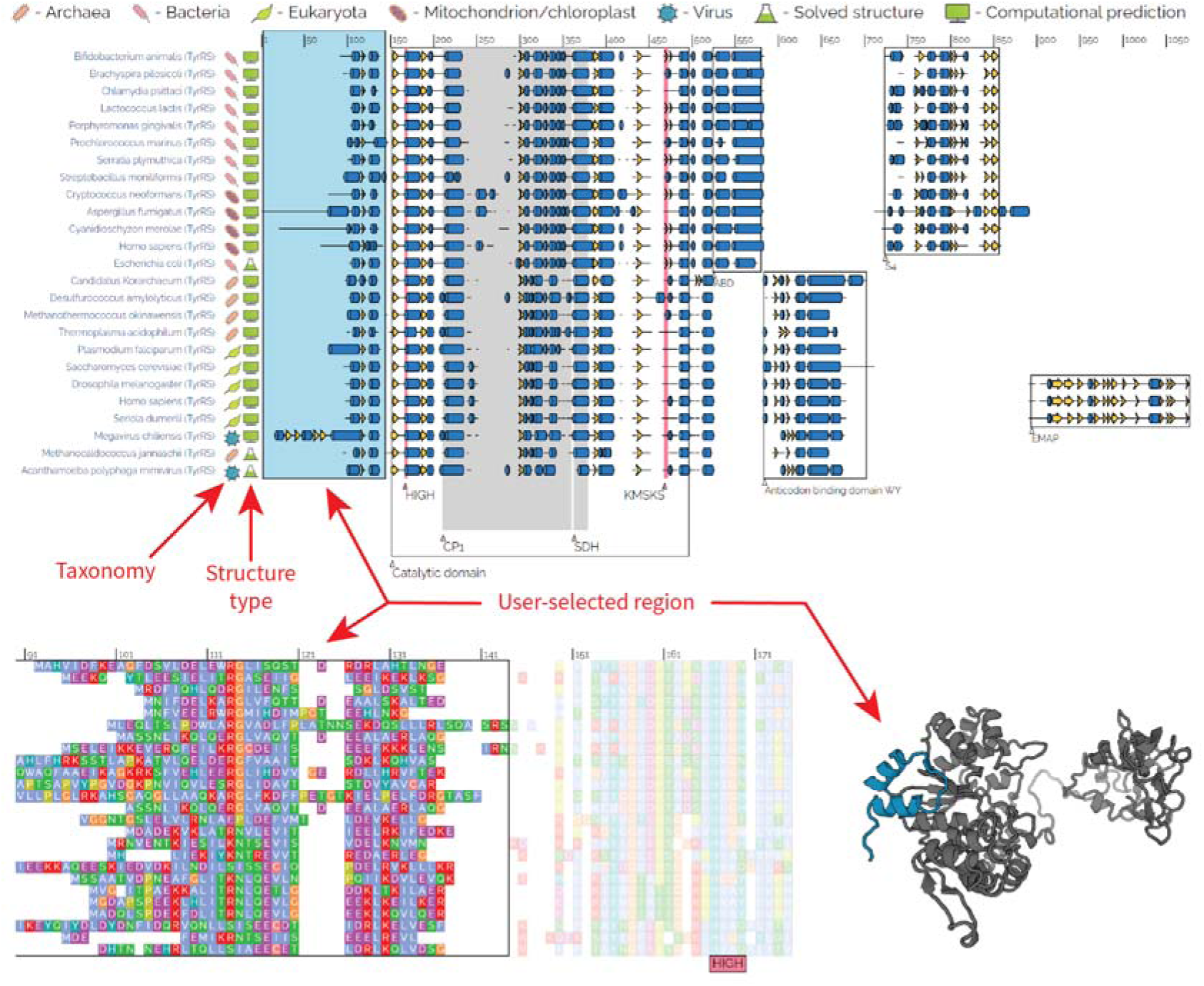
AARS Online display for a multiple sequence/structure alignment of TyrRS. Top: an alignment of domain architectures for TyrRS, HIGH and KMSKS motifs annotated, as identified by Eriani et al. 1990 [127]. Notation - blue cylinders: helices; yellow arrows: - strands, black lines: loops/turns, white space: alignment gaps. Bottom left: the same alignment as above, but zoomed in to show the amino acid sequences. Bottom right: cartoon representation of an AlphaFold prediction for the Homo sapiens cytosolic TyrRS. The sequence and structure are readily navigated by dragging the cursor over this alignment: the user-selected blue rectangle (top) corresponds to the highlighted regions on the primary structure and 3D protein cartoon (bottom).

Most of the enzymes exhibited on AARS Online were assigned putative functions based on evolutionary analysis, without direct experimental validation, and therefore caution is advised during interpretation. We also advise a level of skepticism when interpreting the classification of eukaryotic aaRS into cytosolic and mitochondrial/chloroplastic compartments, or when eukaryotic splice sites have been predicted without support from mRNA sequencing data.

### Related databases

Related databases and services include:

1. 2001: Aminoacyl-tRNA Synthetase Data Bank (AARSDB) [128]. This resource provided annotated aaRS sequence data (at the time of writing, we can no longer access the AARSDB web-server).
2. 2017: Mitochondrial Aminoacyl-tRNA Synthetases & Pathologies (MiSynPat) [21]. MiSynPat documents the disease-causing mutations linked to human mitochondrial aaRS, and much like AARS Online, facilitates ready navigation between sequence and structure. MiSynPat is available at misynpat.org.
3. 2017: Prokaryotic AARS Database [129]. This tool contains a series of aaRS sequence motifs to assist researchers in predicting aaRS functions based on sequence and is available at bioinf.bio.uth.gr/aars.
4. 2019: tRNAviz [116]. tRNAviz provides annotated tRNA structural annotations across the tree of life, allowing the user to identify conserved elements filtered by clade or tRNA isoform. tRNAviz is available at trna.ucsc.edu/tRNAviz/.

All structures on AARS Online are hyperlinked to their sources on GenBank (for AlphaFold structures) or the Protein Data Bank (for experimental structures). The codon table used during translation is specified, according to The Genetic Codes by NCBI,^3^ which was compiled from earlier work [130], [131]. Moreover, aaRS Families are hyperlinked to their corresponding entries in the Prokaryotic AARS Database [129] and MiSynPat [21]. Enzyme commission numbers are used to link the aaRS to the BRENDA enzyme database [132].

### Outlook and limitations

AARS Online provides a friendly introduction to the aaRS and highlights the subtle idiosyncrasies that exist across the tree of life, and across the tree of aaRS. Enzymes are classified into Classes, Subclasses, and Families according to their catalytic domain(s), which is the only structural feature universal to the aaRS. While great care is taken to ensure its accuracy, a healthy level of skepticism is advised when navigating the platform, as many entries in the database are based on bioinformatic predictions: including aaRS functional assignments, gene boundaries, compartmentalization (cytosol or mitochondria/chloroplast), and protein structures (AlphaFold). While experimentally solved structures are generally not faced with the same limitations, they are much fewer in number, often harbor missing residues or truncated domains, and tend to be sourced from a comparatively narrow spectrum of the biosphere. Notwithstanding these caveats, we believe AARS Online to be a useful resource for experts and non-experts alike. With the help of community input, we hope this platform will continue to evolve so that it always reflects the latest research around the aminoacyl-tRNA synthetases.

## Acknowledgements

The authors acknowledge support from the Alfred P. Sloan Foundation Matter-to-Life program Grant number G-2021-16944 (J.D, C.W.C., and P.R.W.), the Canadian Institutes of Health Research and Natural Sciences and Engineering Research Council of Canada (H.C.), the Academy of Finland and Sigrid Juselius Foundation (H.T.), and a Royal Society Newton International Fellowship (C.A.C). Most AlphaFold structures were generated by the New Zealand eScience Infrastructure (NeSI) high performance computing facilities, funded jointly by NeSI’s collaborator institutions and the Ministry of Business, Innovation & Employment’s Research Infrastructure programme.

## Footnotes

Marco Biasini. (2015). pv: v1.8.1. Zenodo. 10.5281/zenodo.20980, available online at https://biasmv.github.io/pv/.

FigTree available at http://tree.bio.ed.ac.uk/software/figtree/

Codon tables are summarized at https://www.ncbi.nlm.nih.gov/Taxonomy/Utils/wprintgc.cgi

